# Phylogeny of docks and sorrels (*Rumex*, Polygonaceae) reveals plasticity of reproductive systems

**DOI:** 10.1101/2020.09.11.293118

**Authors:** Kirstie D. Grant, Daniel Koenemann, Janet Mansaray, Aisha Ahmed, Hamid Khamar, Jalal El Oualidi, Janelle M. Burke

## Abstract

The genus *Rumex* is a unique member of the Polygonaceae (Buckwheat) family of plants. A source of intrigue for *Rumex* lies in the diversity of the reproductive systems associated with the subgenera, species, and subspecies within this genus. Four previously circumscribed subgenera, some 200 species, and a number of subspecies comprise the collective *Rumex* genus. These species exhibit monoecious, dioecious, synoecious (hermaphroditic), and polygamous reproductive systems. Moreover, some of the dioecious species contain sex chromosomes, a phenomenon that is very rare in angiosperms. Apart from the confirmed morphological and phytogeographical distinctions, two of the four described subgenera, *Acetosa* and *Acetosella*, are distinctive in their exhibited sex chromosome systems. For this study, we used three chloroplast markers, *rbcL, trnH*-*psbA, trnL*-*F*, and dense taxon sampling, to reconstruct a molecular phylogeny for *Rumex*.The reconstructed phylogeny for this work resolves six major clades and one large grade in *Rumex*. In addition, the species with known dioecious reproductive systems derived from unique sex chromosome systems are resolved in two different clades nested within “the dioecious clade”. These results suggest that the species with divergent sexual systems are more closely related to each other than to other species comprising the rest of the *Rumex* genus. Furthermore, some species with known synoecious reproductive systems are resolved in a single clade which is also nested within “the dioecious clade”. These results imply a possible reversal occurring over time which suggests the highly plastic nature of reproductive systems among *Rumex* species.

## Introduction

Commonly known as docks and sorrels, *Rumex* L. (Polygonaceae) is a relatively large genus. *Rumex* encompasses four circumscribed subgenera, approximately 200 species, and hundreds of described subspecies or varieties. Many species in *Rumex* are cosmopolitan in nature, spanning six continents of the world. However, many individual species are either regionally endemic, native, or introduced on particular continents (Rechinger, 1937). The cosmopolitan attributes of *Rumex* species are indicative of their ability to thrive in a wide variety of environmental conditions. In addition to being distributed globally, plants of the genus inhabit a wide range of habitats and ecotypes.

*Rumex* species are among the most ubiquitous plants in the world. Described species are just as recurrent in dry and sandy soils as they are in marshes and cultivated fields spanning the arctic, subarctic, boreal, temperate, tropical, and subtropical localities (Löve & Kapoor, 1967). Although several biological species demonstrate little to no niche preference (*e*.*g*., *Rumex crispus, Rumex obtusifolius*), there are others that exhibit exceedingly precise ecological requirements (*e*.*g*., *Rumex bipinnatus, Rumex pictus*). The large variation in the distribution of *Rumex* species might also account for the large deviation observed in the morphology of these species, whereby some reach almost seven meters in height, and others rarely exceed a few centimeters (Rechinger, 1949; Löve & Kapoor, 1967; Rechinger, 1990).

The broad variation in both the morphology and phytogeography of *Rumex* species is also indicative of the substantial taxonomic classification interest in these species. Documented descriptions of plants in the genus date back to the time of classical Greece. Species of *Rumex* are first noted by Hippocrates (greek physician) and Theophrastus (Greek philosopher) under the name *Lapathum* (Campderá, 1819). The first formal monograph of *Rumex* species was completed in 1819 (Campderá, 1819), documenting 110 species of *Rumex* proper and delineating three genera of “Rumices” in the broad sense: *Emex* (L.) Campd., *Rumex*, and *Oxyria* Auth.

Its second formal monograph was completed in 1856 (Meisner, 1856), documenting 134 species of *Rumex*, dividing it into three sections: *Acetosa* (‘sorrels’), *Acetosella* (‘sorrels’), and *Lapathum* (‘docks’) (Meisner, 1856; Löve, 1967). In the 20^th^ Century, progress in the taxonomic and cytological study of *Rumex* was largely accomplished by two researchers: Áskell Löve and Karl Heinz Rechinger (Rechinger, 1937; Rechinger, 1954; Löve, 1967). Löve extensively documented the cytological diversity of *Rumex*, and he proposed a generic status for *Acetosa* and *Acetosella* (the groups with species bearing heteromorphic sex chromsomes) and subgeneric status for *Axillares* and *Platypodium*. Löve also considered *Rumex* to be composed of several smaller genera corresponding to a number of cytotypes (Löve 1957; Löve & Kapoor 1967; Mariotti *et al*., 2006, 2009).

Over the course of his long career, Rechinger effectively monographed *Rumex*, using plant morphology and geographic distribution (Rechinger 1933, 1937, 1939, 1949, 1954a, 1954b, 1984, 1990; Brandbyge & Rechinger, 1989). It was not until the mid-1900’s that Rechinger proposed a subgeneric status for *Platypodium* and maintained *Acetosa, Acetosella*, and *Lapathum* as comparable subgenera (Rechiner, 1954). In important respects, Rechinger’s morphological classification mirrored Löve’s cytological classification. Löve’s cytotypes were largely reflected in Rechinger’s subgeneric sectional system. Rechinger, however, chose to retain *Rumex* as a single genus.

Recent molecular phylogenetic work has sought to resolve the placement of *Rumex* in the Polygonaceae more broadly (Sanchez & Kron, 2008, Sanchez *et al*., 2009; Burke *et al*., 2010; Burke & Sanchez, 2011; Sanchez *et al*., 2011; Schuster *et al*., 2011; Schuster *et al*., 2013; Schuster *et al*., 2015). These studies have placed *Rumex* alongside the other Rumices of Campderá (*Emex* and *Oxyria*), with the addition of *Rheum* as either sister to *Oxyria* (Burke *et al*., 2010; Schuster *et al*., 2011) or to *Rumex* + *Emex* (Schuster *et al*., 2013; Schuster *et al*., 2015). One area that lacks clarity has been the placement of *Emex*, which sometimes appears to be nested within *Rumex* (*e*.*g*., Sanchez *et al*., 2011) and is sometimes placed as sister to *Rumex* (*e*.*g*. Burke *et al*., 2010). The relationships of species within *Rumex*, including the relationship between *Rumex* and *Emex*, continue to be poorly understood due to insufficient sampling and paucity of data. To date, the relationships among species placed within Rechinger’s subgenus *Rumex* are particularly obscure.

The reproductive systems of *Rumex* species vary just as much, or more, than their studied morphologies and geographical distributions. The high degrees of variation in the reproductive systems of *Rumex* species can also speak to the macroevolutionary significance of circumscribed *Rumex* subgenera, another attribute that accounts for much of the longstanding interest in this genus. Species of *Rumex* demonstrate synoecious (hermaphroditic), monoecious, dioecious, and polygamous reproductive systems (Rechinger 1949; Rechinger 1954a; Löve & Kapoor, 1967; Mosyakin, 2005; Navajas-Pérez *et al*., 2005). Most of the reproductive system diversity has been described in subgenera *Acetosa* or *Acetosella*. In particular, most species in those subgenera are dioecious (Rechinger 1937, 1949, 1954a, 1984). A few species in subgenus *Rumex* have variable systems, especially between synoecy and monoecy (*e*.*g*., *Rumex crispus*, pers. obs.). Also noteworthy are the three species of *Rumex* endemic to the Hawaiian islands (*Rumex albescens, R. giganteus* and *R. skottsbergii*), which are all monoecious (Wagner *et al*., 1999).

Heteromorphic sex chromosomes are extraordinarily uncommon in plants, occurring in <1% of all land plants (Ming *et al*., 2011). Chromosomal sex determination systems are restricted to the angiosperms, bryophytes, and gymnosperms, but have evolved repeatedly and independently within these groups (Charlseworth, 2002; Ming *et al*., 2011; Renner, 2014). In plants with sex chromosomes, a variety of sex determining mechanisms are present (Ming *et al*., 2011). In *Rumex*, the documented sex chromosomes are heteromorphic. Two sex determining chromosomal mechanisms are known: XX/XY and XX/XY_1_Y_2_ (Löve, 1940; Löve, 1942; Löve, 1943; Löve, 1944; Löve & Löve, 1948; Shibata *et al*., 1999; Shibata *et al*., 2000; Navajas-Pérez *et al*., 2005; Cunado *et al*, 2007; Ming *et al*. 2011). The XX/XY (male heterogamy) system bears more than a passing resemblance to the mammalian sex determination system, and studies of the formation of this chromosomal arrangement in plants may give insights into the historical formation of the analogous system in mammals (Charlesworth, 2002). The XX/XY_1_Y_2_ system is dosage-dependent, and plant sex is based on the autosome to sex-chromosome ratio. In this system, female individuals have 14 chromosomes, and male individuals have 15 chromosomes (Löve, 1940; Löve, 1944; Löve, & Kapoor,1967; Navajas-Pérez *et al*, 2005).

The vast majority of plants are synoecious with at least morphologically hermaphroditic flowers (Ming *et al*., 2011) and this has been considered the ancestral state for land plants (Navajas-Pérez *et al*., 2005; Ming *et al*., 2011, but see also Renner, 2014). Holding true to the same synoecious ancestral state, the vast majority of *Rumex* species, particularly those nested within subgenus *Rumex*, reveal predominantly synoecious reproductive systems (Rechinger, 1937; Rechinger, 1954; Świetlińska, 1963). Dioecious plants have been proposed to be derived from synoecious ancestors via two possible pathways, with either gynodioecy as a transitionary state or monoecy as a transitionary state (Figure 1; Charlesworth & Charlesworth, 1978; Lewis,1942; Lloyd, 1980; Lloyd & Webb, 1986; Renner & Won, 2001; Barrett, 2013; Crossman & Charlesworth, 2013).

**Figure 1.**
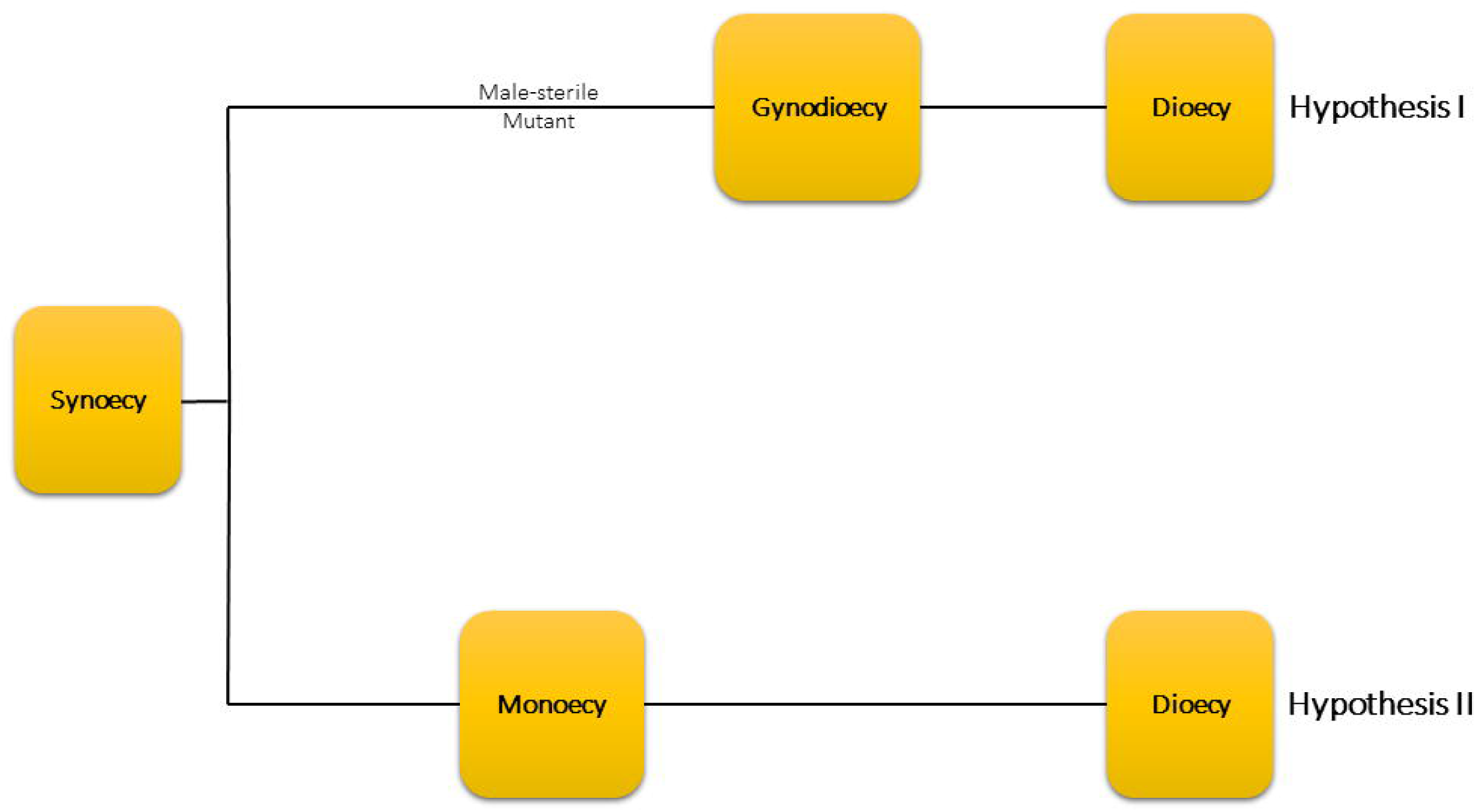
Hypothesis I proposed gynodioecy as the most likely intermediary state in the pathway to dioecy (Charlesworth & Charlesworth, 1978). Hypothesis II proposed monoecy as the most likely intermediary state in the pathway to dioecy (Lewis, 1942).

Both monoecy and dioecy have evolved multiple times in plants (Charlesworth 2002, Renner & Won 2001; Ming *et al*., 2011). Historically, it had been suggested that dioecy emerged multiple times in *Rumex* (Fox, 1985). Navajas-Pérez *et al*. (2005), however, concluded that heteromorphic sex chromosomes evolved only once in *Rumex*, while dioecy (without heteromorphic sex chromosomes) evolved twice. For some time, dioecy appeared to be evolutionarily terminal since many dioecious taxa are embedded in clades of hermaphroditic taxa and purely dioecious clades are usually species-poor (Heilbuth, 2000; Barrett, 2013).

More recently, however, several cases of reversals from dioecy to monoecy have been recorded (Barrett, 2013). In all of these cases, monoecy was derived from a group of “leaky” dioecious plants. At the population level of these supposedly dioecious taxa, some hermaphroditic and monoecious individuals are always present (Świetlińska, 1963; Lloyd 1980, Charlesworth, 2002; Barrett, 2013; Crossman & Charlesworth, 2013). The exact conditions under which dioecy, especially dioecy controlled by sex chromosomes, transitions back to monoecy or synoecy are unknown and only a few cases have been well studied (Heilbuth, 2000; Schaefer & Renner, 2011; Crossman & Charlesworth, 2013).

The purpose of this study was to provide a molecular phylogeny of *Rumex*, test the placement and monophyly of its circumscribed subgenera, and to elucidate the evolution of reproductive systems in *Rumex*. We here present a new phylogeny of *Rumex* constructed using three plastid gene regions (*trnH-psbA, rbcL*, and *trnL*-*F*) and 67 *Rumex* species. One objective is to discover whether Rechinger’s subgeneric delineations based on morphology and phytogeography are supported by our phylogeny based on molecular data. In addition, we address whether large scale patterns can be discerned in the reproductive systems exhibited by *Rumex*.

## Materials and Methods

### Taxon Sampling and DNA Isolation

DNA was isolated from 109 accessions, representing 67 *Rumex* species. Of the 109 included accessions, a total of 99 *Rumex* accessions, 6 *Rheum* species, 3 *Emex* accessions, and 1 species of *Persicaria* are represented. *Persicaria virginiana, Rheum alexandrae, Rheum emodii, Rheum nobile, Rheum officinarumas, Rheum palmatum*, and *Rheum rhabarbarum*, were included as outgroup species. Additional plant samples were obtained through the GenBank sequence database (Appendix 1). Samples were taken from a combination of herbarium specimens (K, NY, OSC, RAB, US), field collections, and cultivated samples from collaborators. Herbarium acronyms follow the Index Herbariorum (Thiers, 2019).

All fresh leaf samples were dried using silica gel. Plant tissue was homogenized using the FastPrep-24™ 5G Sample Preparation System (M. P. Biomedicals, LLC Santa Ana CA, USA). Total genomic DNA was extracted from herbarium specimen-sampled and silica-dried leaf tissues using a BIOLINE ISOLATE II Plant DNA Kit (Cat No. BIO-52070). Modification for herbarium material proceeded as follows: Cell lysis was carried out using 300µL of buffer (PA1 or PA2) and 30µL of proteinase K (20µg/mL) and incubated for 18 hours at 65□ on an orbital shaker).

### Marker Selection

For this first comprehensive phylogeny for the genus, we focused on plastid marker selection. Previous authors of recently reconstructed Polygonaceae phylogenies have used nrITS as a nuclear marker (Schuster *et al*., 2011; Schuster *et al*., 2015). However, we avoided nrITS for this phylogeny due to a number of issues that would interfere with accurate reconstruction of evolutionary relationships: 1) nrITS is extremely variable and difficult to align (66% of nrITS sequence data was excluded in Schuster *et al*. (2015) publication, and 2) Due to widespread polyploidy documented in multiple *Rumex* species, sequences of nrITS would not necessarily be low copy, and there would be substantial issues with paralogy and orthology across multiple polyploidy events.

For plastid marker selection, we screened multiple markers that had previously been used in Polygonaceae reconstruction (Burke *et al*., 2010; Burke and Sanchez, 2011; Koenemann and Burke, 2020). We selected markers that both showed sufficient variation across the genus, and were easily amplified for most taxa.

### PCR Amplification and Sequencing

Amplification of DNA markers was completed for three plastid regions: *rbcL, trnH-psbA* and *trnL-F*. The first amplified region was the plastid large subunit of ribulose-bisphosphate carboxylase (*rbcL*) using the primers *rbcLaF* (5□-ATG TCA CCA CAA ACA GAG ACT AAA GC-3□) and *rbcLaR* (5□-GTA AAA TCA AGT CCA CCR CG-3□) (Table 1). PCR conditions were as follows: 94□ for 1 min, followed by 34 cycles of 94□/15 s, 54□/15 s, and 72□/30 s, and a final extension period of 5 min at 72□. The second region was analyzed using primers *trnH* (5’-ACT GCC TTG ATC CAC TTG GC-3’) and *psbA* (5’-CGA AGC TCC ATC TAC AAA TGG-3’) as an intergenic spacer. PCR conditions were as follows: 94□ for 2 min, followed by 34 cycles of 94□/30 s, 55□/30 s, and 72□/30 s, and a final extension period of 7 min at 72□. Compared to the *rbcL* gene and the *trnH-psbA* intergenic spacer, a second, much shorter intergenic spacer was examined. This intergenic spacer was amplified using primers *3’trnL*^*UAA*^*F* (5’-GGT TCA AGT CCC TCT ATC CC-3’) exon and the *trnF*^*GAA*^ (5’-ATT TGA ACT GGT GAC ACG AG-3’) gene which were used as primers. PCR conditions were as follows: 80□ for 5 min, followed by 34 cycles of 94□/1 min, 55□/1 min, and 72□/2 min, and a final extension period of 5 min at 72□. PCR was performed to amplify all target gene regions using a BIOLINE MyTaq™ Red Mix, 2X (Cat. No. BIO-25044) with no special PCR conditions. PCR samples were then visualized on 1% agarose gels and run at 100V for 15 - 30 min against the BIOLINE 100bp – 2000bp EasyLadder I (Cat. No. BIO-33045) to observe the bands of specified gene regions. PCR experiments were segregated to contain amplification of only fresh or only herbarium material to help prevent cross-contamination.

**Table 1.**
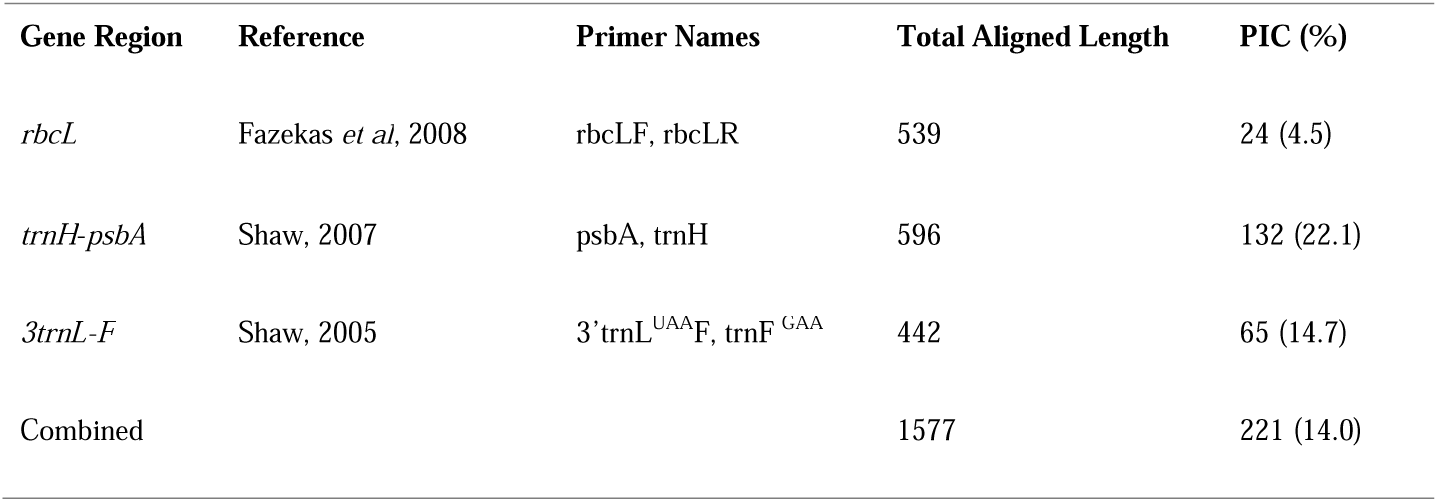
Gene regions used: name of primers, total length of region, % parsimony informative characters

PCR amplicons were sent to Eurofins Genomics (Louisville, KY) for Sanger sequencing. Sequences were edited using Geneious v. 10 (Biomatters Ltd.). Reviewed sequences were aligned with MUSCLE (Edgar, 2004), and concatenated using MESQUITE (Maddison, 2005).

### Phylogeny Reconstruction

The final dataset contained 92 *rbcL*, 93 *trnL-F*, and 95 *trnH-psbA* accessions (Grant, 2020). All phylogenetic analyses were completed using the CIPRES Science Gateway V 3.3 (Miller *et al*., 2010). Prior to the phylogenetic reconstructions, we performed ModelTest-NG (Darriba *et al*., 2019) for the concatenated matrix to determine the suggested model of evolution. ModelTest-NG indicated that the best fit was the General time reversible (GTR) model.

We performed Maximum likelihood (ML) phylogeny reconstruction using GARLI v. 2.01.1067 (Zwickl, 2006). We used the default GARLI parameters with the following exceptions. We performed 1000 search replications (10 iterations of 100 search replicates). In order to better search tree space, we increased the attachments per taxon setting to 150 and extended the generations without improvement parameter to 50000. To evaluate support for phylogenetic relationships, statistical bootstrapping was performed, specifying only one search replicate per bootstrap iteration for 100 iterations. All bootstrap trees were downloaded and used to generate a majority rule consensus tree in MESQUITE (Maddison, 2005). The consensus tree was visualized in FigTree version 1.4.3 (Rambaut, 2014).

We performed Bayesian Inference phylogeny reconstruction in MrBayes 3.2.7a (Ronquist *et al*., 2012). The priors were set to the defaults (Dirichlet). We set the seed number at 123. We conducted two independent Markov Chain Monte Carlo (MCMC) runs, each with four chains employing BEAGLE library acceleration (as recommended by CIPRES). Each MCMC run was set to complete 5 million generations, with trees sampled every 1,000 generations. The first 25% of trees in each run were discarded as burn-in. MrBayes then synthesized the two independent runs and we extracted the majority rule consensus tree with posterior probabilities.

Posterior probability and bootstrap values were visualized using FigTree version 1.4.3 (Rambaut, 2014) and MESQUITE (Maddison, 2005). Posterior probabilities above 90% and bootstrap support values above 70% were considered significant and annotated in the final phylogeny.

## Results

The recovered most likely tree was generated using 109 specimen accessions. This included 7 outgroup species, 3 accessions of *Emex*, and 99 accessions of *Rumex*. The present phylogeny represents 67 *Rumex* species, more than twice the number of species of *Rumex* sampled in previous phylogenies (31 species in Navajas-Pérez *et al*., 2005; 13 species in Schuster *et al*., 2015). A total of 47 sequences were missing from the final matrix, yielding 14.4% missing data in the final analysis (Grant, 2020). Table 1 summarizes the variability of each of the gene regions. The most variable region was *trnH-psbA*, which consisted of 22.1% parsimony informative characters. The least variable region was *rbcL* which consisted of 4.5% parsimony informative characters. The most likely tree recovered by GARLI received a likelihood score of Ln= -5767.548440.

The genus *Rumex* was recovered as monophyletic with strong support (100 Bayesian Posterior Probability/98 Maximum Likelihood Bootstrap) (Figure 2). The analysis did not recover *Rumex* subgenus *Rumex*, the subgenus with the most species diversity, as monophyletic. In our phylogeny, species of subgenus *Rumex* form a grade at the base of the tree (“Basal Grade” - Figure 2). Above the *Rumex* grade, *Emex* (Clade 1), was recovered as monophyletic and sister to “the dioecious clade” (Figure 2). While the results indicate strong support for the relationship between the known *Emex* species, *E. australis* and *E. spinosa* (100/98), they are conflicting and show poor support for the placement of *Emex* within *Rumex*. Posterior probability support for the placement of *Emex* as sister to the Dioecious Clade is only 52% and the most likely GARLI tree placed *Emex* within the Basal Grade of subgenus *Rumex*. Furthermore, different gene regions reconstructed conflicting topologies for the placement of *Emex*. The *rbcL* phlyogeny placed *Emex* within *Rumex* subgenus *Rumex* (50% bootstrap support). Both *trnh-psbA* and *trnL*-*F* placed *Emex* as sister to the *Rumex* genus (*trnh-psbA* <50% bootstrap support, and *trnL*-*F* 91% bootstrap support) (results not shown).

**Figure 2.**
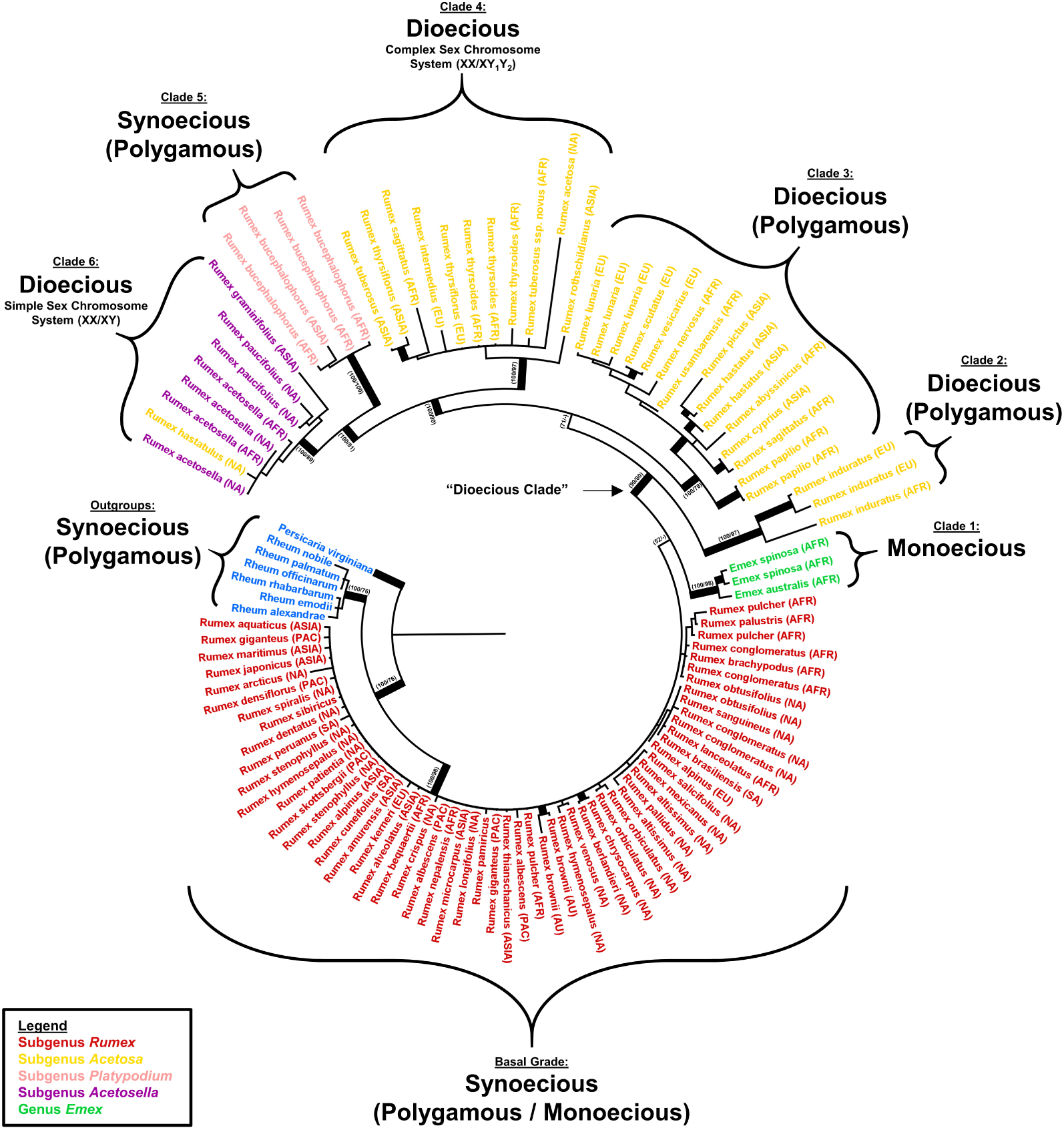
Bayesian phylogenetic reconstruction for *Rumex* species using three chloroplast sequences (*rbcL, trnH*-*psbA*, and *trnL*-*F*). Thickened branch indicates simultaneous posterior probability above 90% and bootstrap support above 70%. Exact support values are indicated at important nodes (Bayesian Posterior Probability / Maximum Likelihood Bootstrap). Outgroup species (*Rheum* and *Persicaria*) are shown in blue. *Rumex* species traditionally placed in subgenus *Rumex* are shown in red. Species traditionally placed in the sister genus *Emex* are shown in green. *Rumex* species traditionally placed in subgenus *Acetosa* are shown in gold. *Rumex* species traditionally placed in subgenus *Platypodium* are shown in pink. *Rumex* species traditionally placed in subgenus *Acetosella* are shown in purple. The arrow denotes the “Dioecious Clade” referenced in the text. Parenthetical abbreviations following the taxa represent collection localities: NA = North America, ASIA = Asia, AFR = Africa, EU = Europe, PAC = Pacific, SA = South America, AU = Australia. Accessions that lack locality information were composed of GenBank sequences, where collection locality could not be determined.

The remaining taxa, comprising the subgenera *Acetosa, Acetosella*, and *Platypodium* form a highly supported (99/80) monophyletic group (Figure 2). This group is denoted as “the dioecious clade” because the known dioecious *Rumex* species were resolved in this group. The relationships of the clades within this group are also well-supported. Our recovered phylogenetic tree did not recover subgenus *Acetosa* as monophyletic. Within the dioecious clade, subgenus *Acetosa* is comprised of three well-supported, monophyletic groups, Clade 2 (100/97), Clade 3 (100/78), and Clade 4 (100/97), and is nested below a pair of clades, represented by subgenus *Platypodium* (Clade 5) and subgenus *Acetosella* (Clade 6). The pair is also well supported (100/81). Subgenus *Platypoidium* was recovered as monophyletic with strong support (100/100), and consists of four accessions of its only circumscribed species: *Rumex bucephalophorus*. Species in subgenus *Acetosella* were recovered together with strong support (100/89), but the inclusion of *Rumex hastatulus* means the subgenus was not recovered as monophyletic (Figure 2).

Beginning at its basal lineages, the recovered topology largely corresponds to the diversity of the reproductive and sex chromosome systems present in *Rumex*. Species in subgenus *Rumex* are hermaphroditic with no documented heteromorphic sex chromosomes. These species, while not recovered as a clade, are recovered together in the basal grade. Also with no documented heteromorphic sex chromosomes, *Emex* is represented as a clade that consists of purely monoecious species.Within “the dioecious clade”, subgenus *Acetosa* consists entirely of dioecious species, with some members exhibiting the sex chromosome system XX/XY_1_Y_2_ represented in Clade 4 (Dioecious Species, Complex Sex Chromosome System) (Figure 2). Subgenus *Platypodium*, another hermaphroditic group with no reported sex chromosomes is nested between subgenera *Aceotsa* and *Acetosella*. Subgenus *Acetosella* (Clade 6) consists of species that are both dioecious and have the sex chromosome system XX/XY (Figure 2).

## Discussion

Our results produced a phylogeny of *Rumex*, with six clades and one grade, largely congruent with Rechinger’s subgeneric classification and the evolution of reproductive and sex chromosome systems present in *Rumex*. Moreover, the species with divergent sex chromsome systems in *Rumex* are resolved as two separate clades, with the simple sex chromosome system, XX/XY, being more derived. Where the basal grade contains most of the synoecious species, the larger clade in the phylogeny, designated “the dioecious clade”, represents most of the diversity and variation across reproductive systems in the genus.

Within the phylogeny: the basal grade is mostly made up of synoecious species from *Rumex* subgenus *Rumex*. In this portion of the study, we observed a peculiar finding where subgenus *Rumex* was recovered as a grade instead of the anticipated clade. Given this finding, we suspect that potentially, with even more taxon sampling, subgenus *Rumex* would have been recovered as a monophyletic clade. Although dioecious, the species included in Clade 2 and Clade 3 have no reported sex chromosome systems. The species included in Clade 4 exhibit a complex sex chromosome system (XX/XY_1_Y_2_). This placement suggests that this heteromorphic sex chromosome system was derived from dioecious ancestors. The genetic origin of hetermorphic sex chromsomes in *Rumex* is beyond the scope of this manuscript, but we provide a framework to investigate potentially intermediary taxa that may contain homomorphic or transitionary sex chromosome systems.

Subgenus *Platypodium* (Clade 5) was resolved as monophyletic and nested within “the dioecious clade”. Based on its plant and chromosome morphology, earlier studies concerning *Rumex bucephalophorus* have referred to it as the link between subgenus *Rumex*, which is predominantly synoecious, and subgenus *Acetosella*, which is predominantly dioecious (Löve, 1944). Although morphologically variable, *R. bucephalophorus* consistently exhibits a synoecious reproductive system. Its derivation from among the dioecious species in this phylogeny suggests a reversal from a dioecious condition.

The appearance of *R. bucephalophorus* within “the dioecious clade” can also speak to (1) the high degree of reproductive system plasticity and (2) the possibility of a reversal from dioecy to hermaphroditism. This is significant, because it goes against the evolutionary theory which suggests that dioecy may be an evolutionary ‘dead-end’ (Heilbuth, 2000). Reversals from dioecy back to hermaphroditism that typically occur at or near the onset of sex chromosome fluctuations have been documented in some plants (Ming, 2011; Bachtrog *et al*., 2014). This finding is one that would support the placement of *R. bucephalophorus* as it is in the current phylogeny, between subgenera *Acetosa* (XX/XY_1_Y_2_) and *Acetosella* (XX/XY).

Subgenus *Acetosella* (Clade 6), was not recovered as monophyletic. Known dioecious species, *R. hastatulus*, of subgenus *Acetosa* is nested within subgenus *Acetosella. Rumex hastatulus* is documented to exhibit two chromosomal races: a complex sex chromosome system (XX/XY_1_Y_2_, North Carolina Race) which is characteristic of subgenus *Acetosa* and the simple sex chromosome system (XX/XY, Texas Race) which is characteristic of subgenus *Acetosella* (Navajas-Pérez *et al*., 2005; Mariotti *et al*., 2009; Hough *et al*., 2014). In addition, Rechinger’s 1937 treatment indicates a polygamous reproductive system for *R. hastatulus* (Rechinger, 1937). Given the variability found within this speceis, *R. hastatulus* could have theoretically evolved in either subgenus (*Acetosa* or *Acetosella*), where species appear to have diversified according to the type of sex chromosome system they exhibit. Again, this speaks to the plasticity of reproductive and sex chromosome systems within *Rumex*, as a single species can exhibit two different chromosomal ‘races’.

In all, this work has provided a reconstructed phylogeny that differs from those currently published (Navajas-Pérez *et al*., 2012; Schuster *et al*., 2015) and tests the placement and monophyly of its circumscribed subgenera. Additionally, this work has begun to elucidate the evolution of reproductive systems in *Rumex* by way of its proposed schematic for the pathway from synoecy (hermaphroditism) to dioecy. Simultaneously, the reconstructed phylogeny emphasizes the high degree of reproductive system plasticity of *Rumex* species. Where lacking, we have increased taxon density which has given rise to a more comprehensive evolutionary history of *Rumex* where the taxa are concerned. Future directions in *Rumex* research include the identification and application of nuclear markers that will allow for a more robust phylogeny to increase the strength of support for molecular inferencies.

## Acknowledgements

This study was largely funded by award NSF-HRD #1601031 to JMB. The authors would like to thank staff at the following herbaria for faciliating access to specimens for sampling: K, NY, OSC, RAB, and US. Special thanks to Spencer Barrett and Joanna Rifkin for access to cultivated *Rumex* material. We also thank Daniel Atha for providing us with a record of his *Rumex* collections, and faciliating access to silica-dried material. Special thanks to Mr. Ibrahim El Hafid for his assistance with transportation during the Morocco expedition. We also thank Ms. Maria Ramos for her assistance with plant cultivation at the Howard University greenhouse.

## Appendix 1

List of taxa sampled, GenBank Accession Numbers, and Sequence Vouchers Parenthetical values following the voucher are institutional barcodes or accession numbers when available. Chloroplast region order is *rbcLa, trnH*-*psbA*, and *3trnL*-*F* unless otherwise indicated. For sequences that we did not generate, accession information is given as found on GenBank.

### GenBank Sequences Used for this Study

*rbcL*: *Rumex pamiricus* Rech. f. - JF944139.1, *Rumex sibiricus* Hulten-KC483892.1

*trnH-psbA*: *Rumex pamiricus*-JN047053.1

### DNA Sequences Generated for this Study

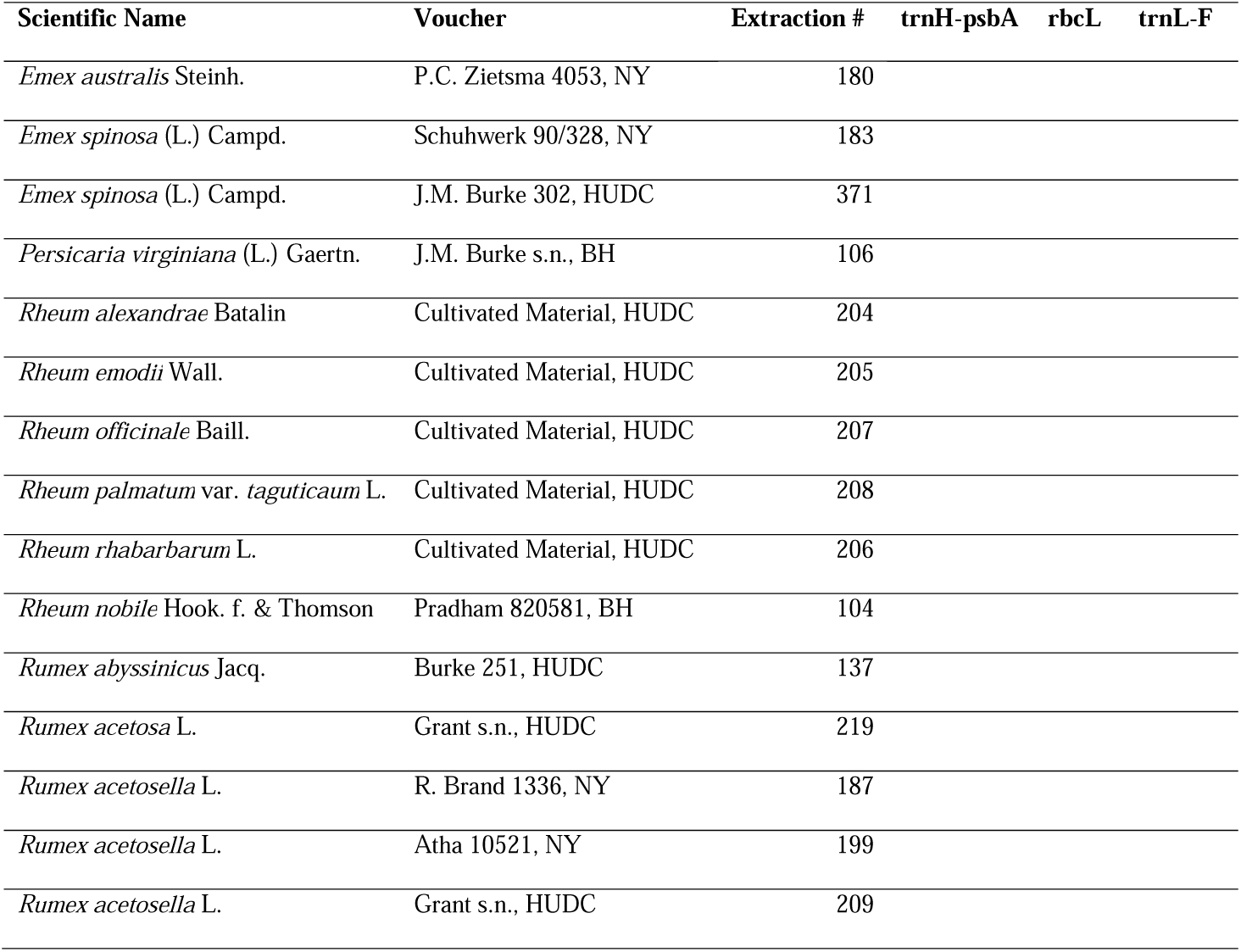
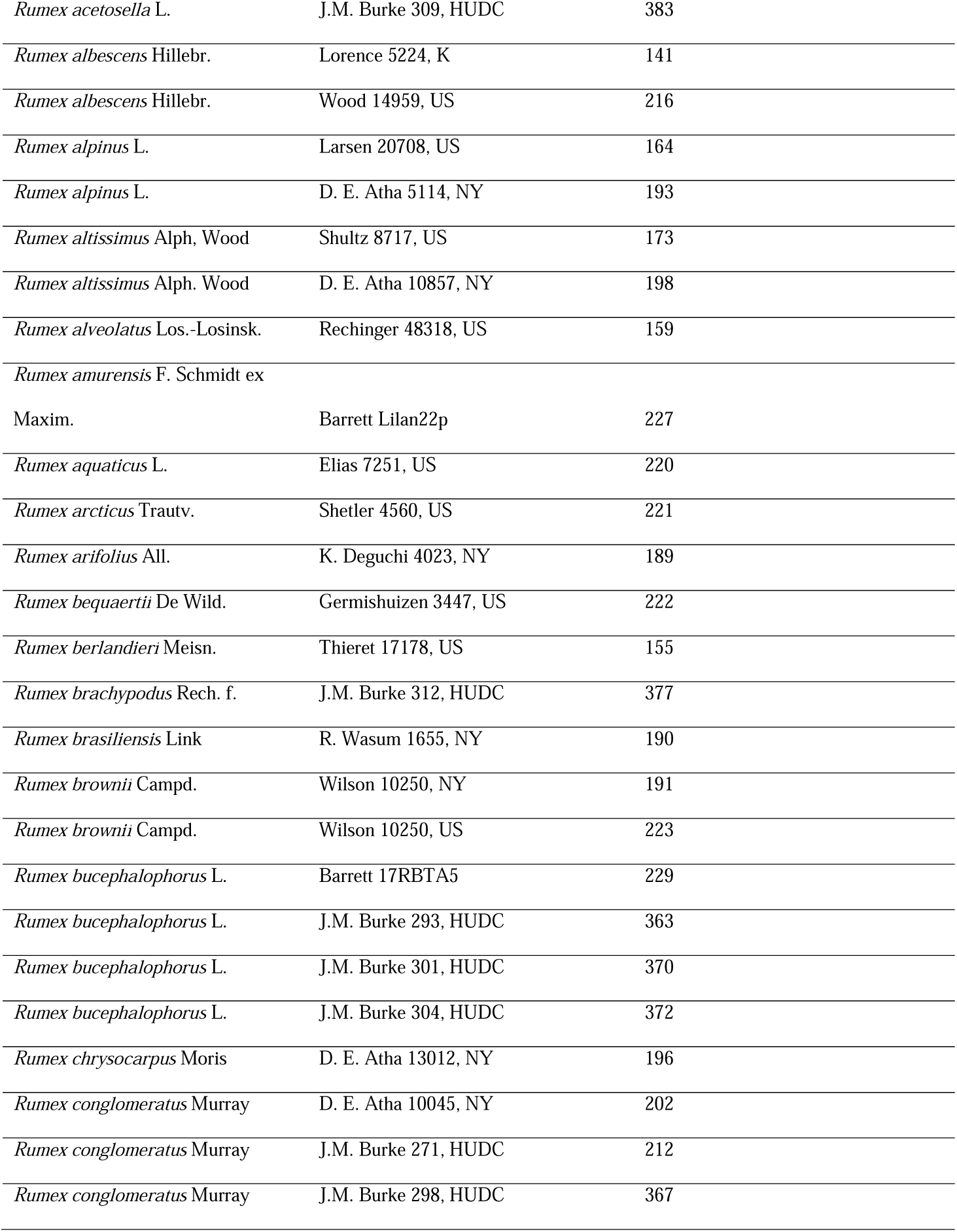
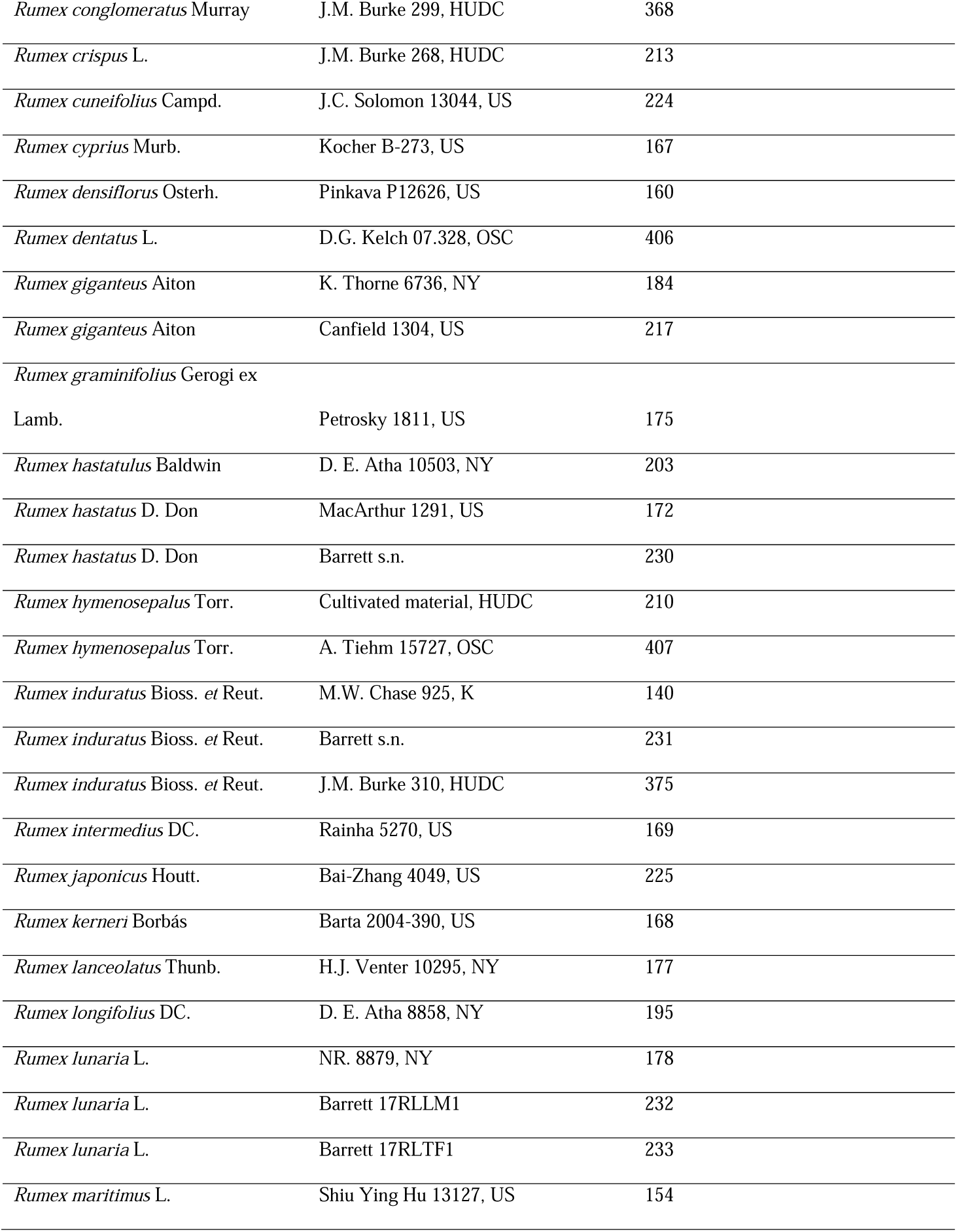
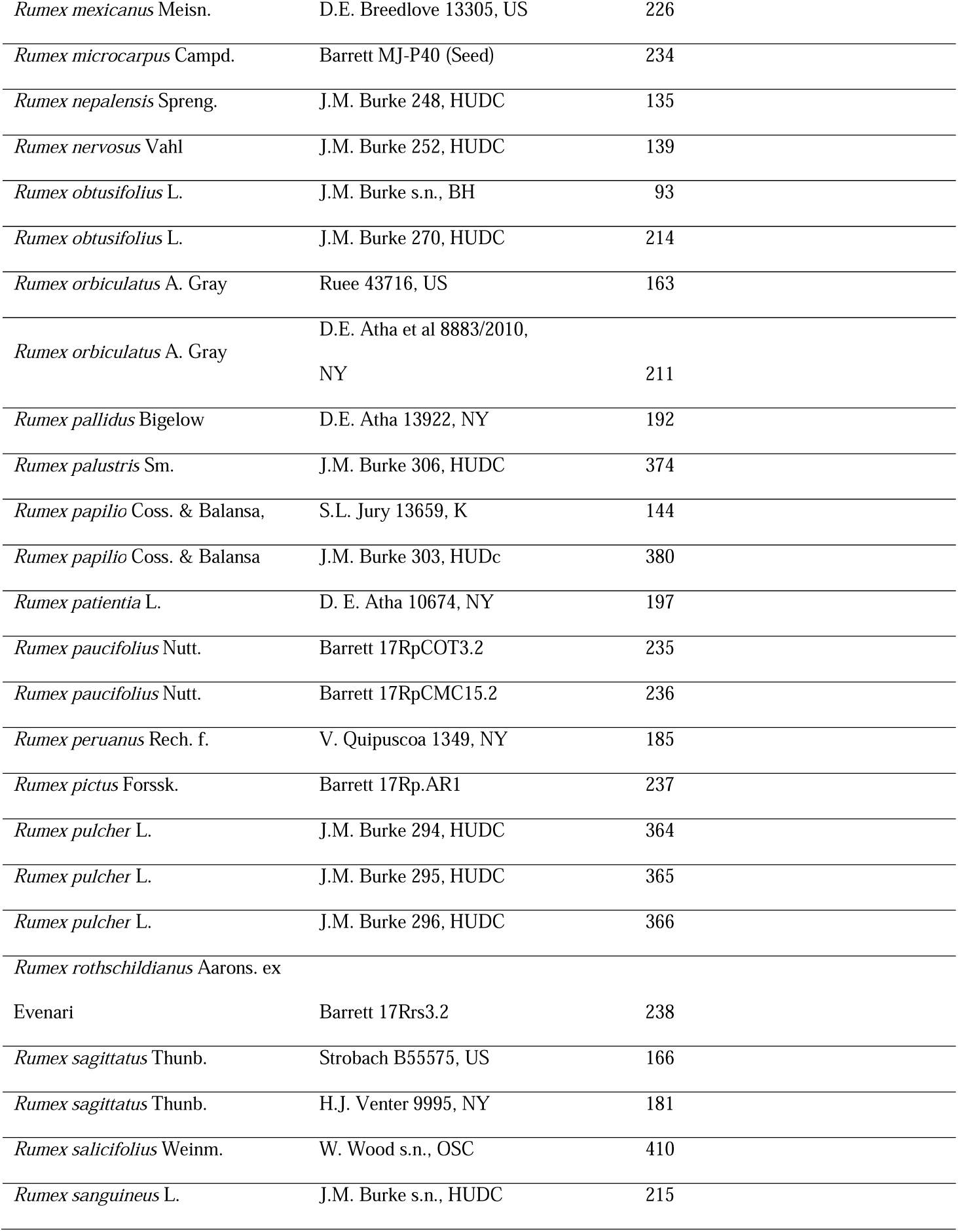
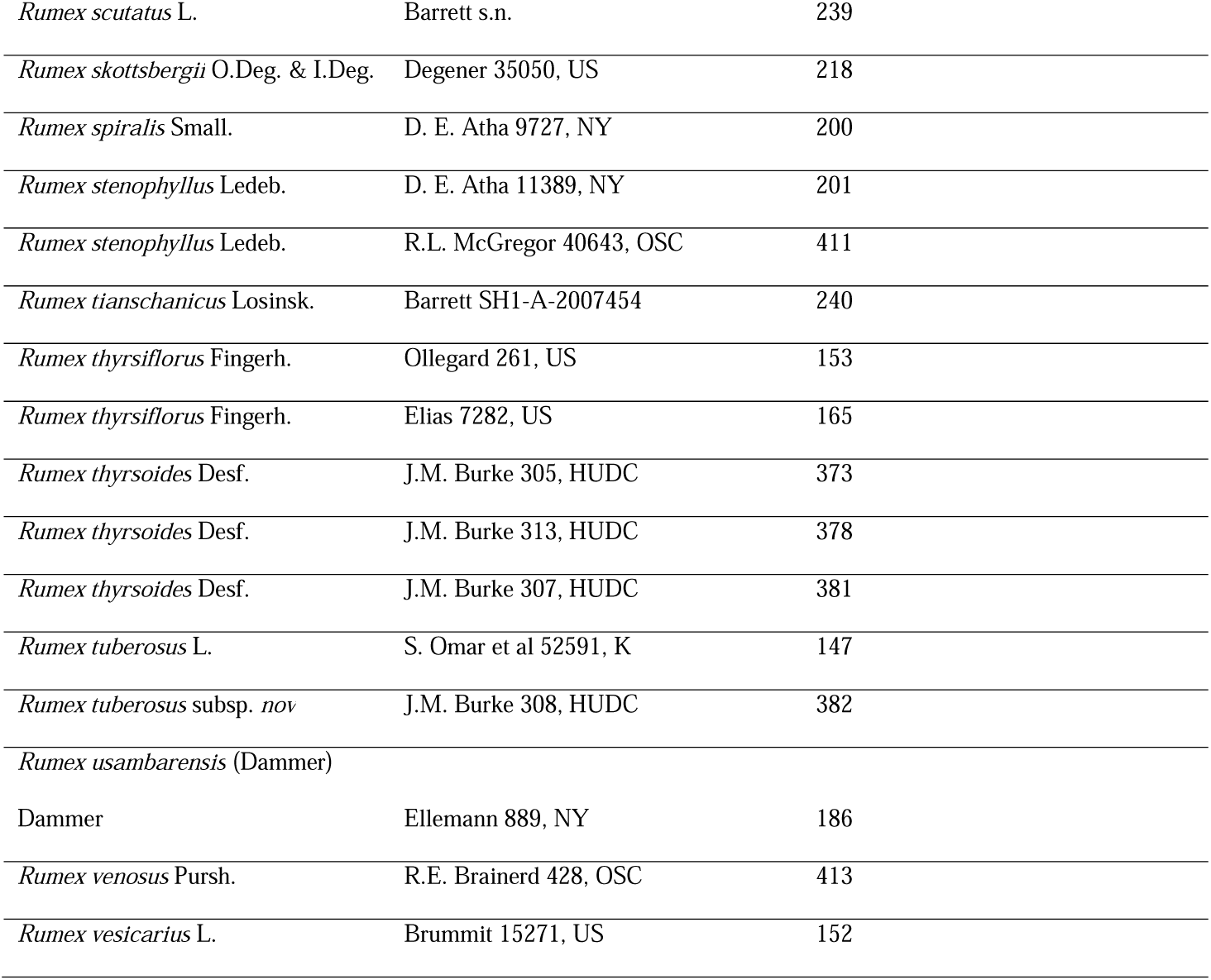

